# Intracellular photoswitchable neuropharmacology driven by luminescence from upconverting nanoparticles

**DOI:** 10.1101/2019.12.17.865261

**Authors:** Jun Zhao, Graham C. R. Ellis-Davies

## Abstract

Photoswitchable drugs are small-molecule optical probes that enable chromatically selective control of drug efficacy. Such light-driven neuropharmacology normally uses UV-visible light. Here we report that luminescence from a NaYF_4_:TmYb nanoparticle can be used for “remote control” of the configuration of an azobenzene-based quaternary ammonium photochrome called “AAQ”. Normally the thermodynamically favored *trans* configuration of AAQ blocks voltage-gated potassium channels. Such activity is reduced by UV irradiation, due to the photochemical *trans* to *cis* isomerization generated by UV light. Since *cis*-AAQ absorbs more blue-green light, this wavelength range can be used to reverse the effects of UV light. We found that in place of such direct photostimulation, the blue luminescence from NaYF_4_:TmYb upconverting nanoparticles could drive AAQ activation inside living cells so as to enable bi-directional control of voltage-gated ion channels using UV and near-infrared light.

---

Photochemical switches are chromophores that exist in two structural configurations which absorb different wavelengths of light^1^. In a biological context the azobenzene (AB) chromophore has been deployed in photoswitches for the control of numerous processes since the 1960s^2,3^. In 1971 Erlanger and co-workers showed that AB derivatives could be used to “photosensitize” membrane bound ion channels, in their case nicotinic acetylcholine receptors, with photoswitchable quaternary ammonium ions^4^. More detailed studies of the biology of their probes appeared in 1980/2, in collaboration with Lester and co-workers^5,6^. Recently, this approach was revived by a set of three groups at Berkeley, who developed rationally targeted quaternary ammonium AB probes in 2004^7^, followed by freely diffusible photoprobes in 2009^8^. Using *cis*-*trans* isomerization, the latter block voltage-gated ion channels in neurons^8^.

Functionality relevant for this biological activity is connected the AB core via *para* di-amido substituents, with both symmetrical or asymmetrical probes producing a use-dependent block of voltage-gated cation channels^8^. In all cases the *trans* configuration is more stable thermodynamically, and absorbs UV light much more strongly than the *cis* configuration, with the latter absorbing slightly more visible light (range ca. 450-530 nm)^2^. However, there is almost complete overlap of the absorption spectra of both configurations^7,8^. The basic structure-activity relationship (SAR) of these photoswitchable drugs is well defined, with the more extended *trans* configuration able to bind more tightly to inside of ion channels in comparison to the more bulky *cis* configuration^8^ (Fig. 1).

**Figure 1.**
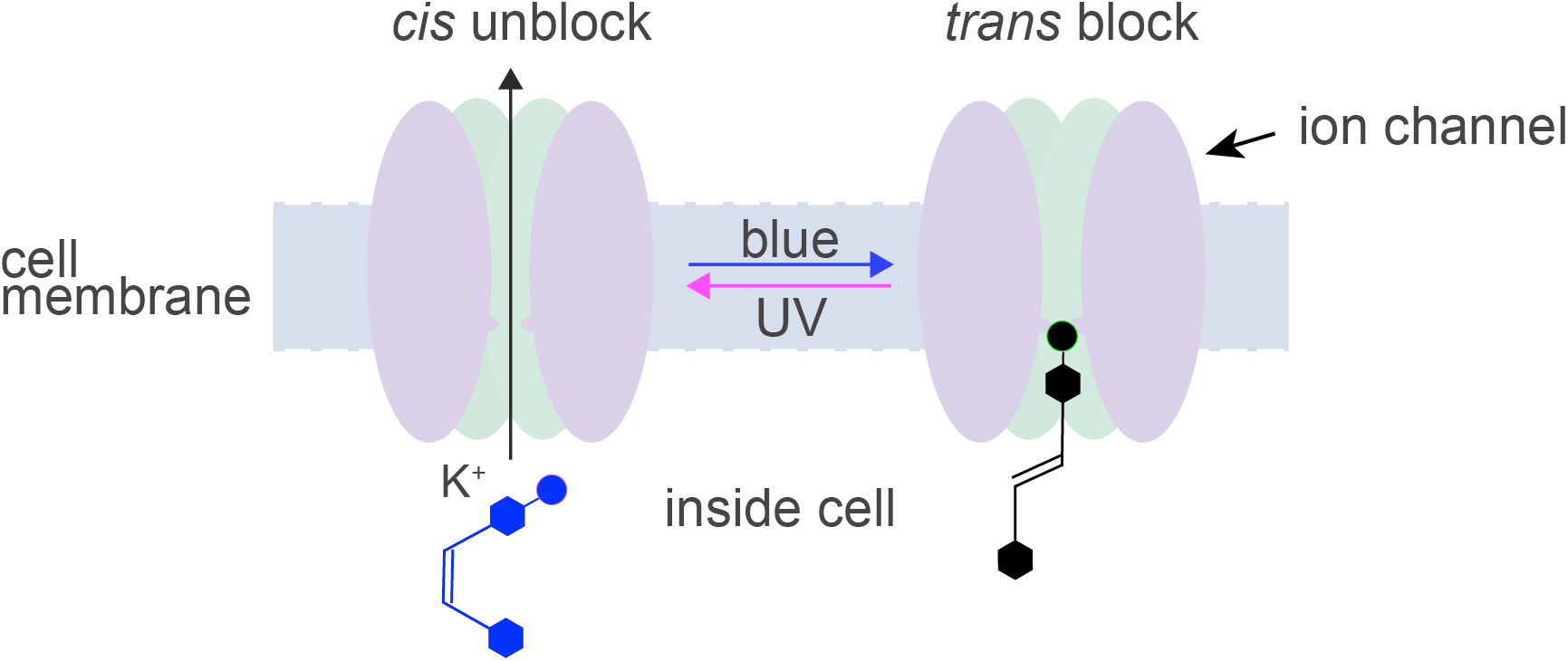
Cartoon of ion channel block-unblock with a photoswitchable drug. Schematic illustration of how *trans* configurations enter into the inner vestibule of membrane-straddling voltage-gated ion channels. Irradiation of the *trans* form with UV light (ca. 365 nm) drives the drug into its *cis* configuration which dissociates from the ion channel, allowing conduction to take place.

Beyond this simple AB core photochrome, much effort has been expended to develop ABs that absorb at longer wavelengths, with orange/red absorbing molecules being something of a “Holy Grail”^9–11^. While much success has been reported in this regard^11^, there seems to be a serious cost-benefit in terms of the drug SAR, with the substituents required for bathochromic shift having deleterious consequences in terms of efficacy, and even solubility^12^. An alternative to the direct absorption of orange light to create the first excited singlet state (S_1_), is two-photon absorption of near-infrared light (NIR). In this modality, excitation to S_1_ Of the AB goes via an ultra-short-lived virtual state (10^−17^ s) due to the simultaneous absorption of two NIR photons^13^. An attractive alternative to such direct NIR excitation is to partner^14–17^ organic chromophores with upconverting nanoparticles (UCNPs) that consist of rare earth elements such as Yb with Tm, Er, Y, Ho, etc. Such materials undergo absorption of single NIR photons by a Yb sensitizer which has a suitable resonance with activator partners (Tm, Er, Y, Ho) to enable facile multiphoton transfer to the latter (Fig. 2). Real excited states of rare earth elements are involved that have much longer life times that the virtual state that is intermediate to S_0_ - S_1_ transitions^14^. Radiative decay of the sensitizers occurs in the visible range, i.e. at shorter wavelengths that the NIR excitation. Thus, the sequential absorption and transfer of multiple NIR photons gives an anti-Stokes shifted emission and, hence, is called “upconversion”.

**Figure 2.**
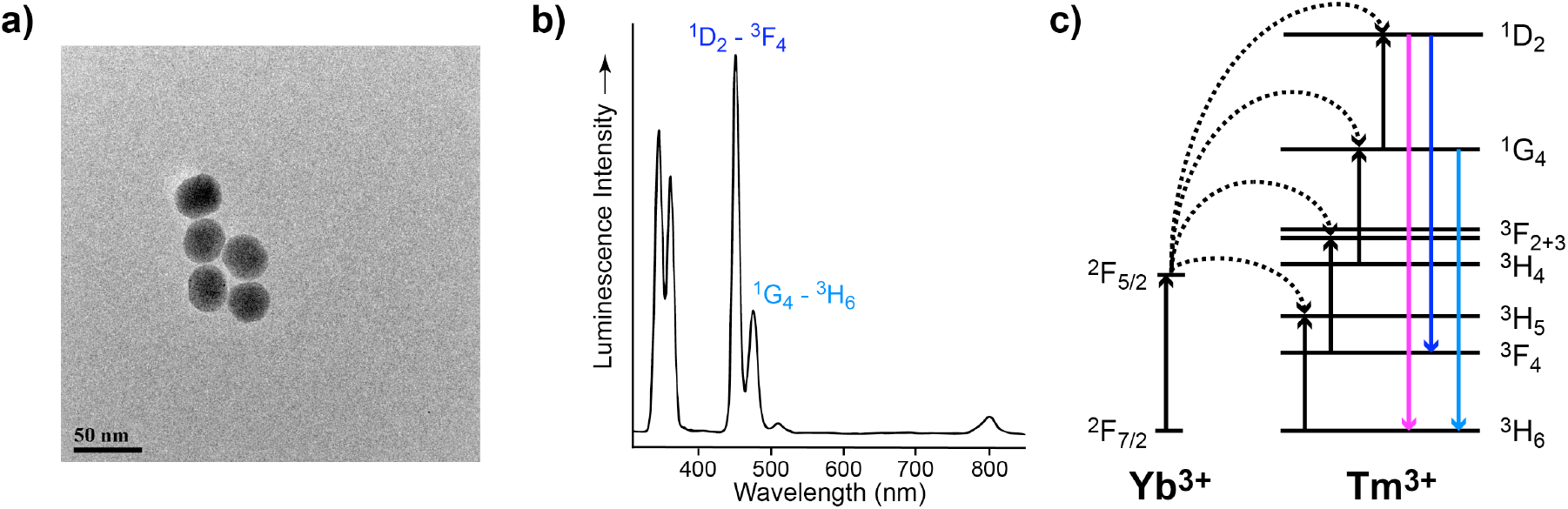
Image of NaYF_4_:TmYb nanoparticles and spectral output. (a) Transmission electron micrograph image of the UCNPs. (b) Luminescence spectrum of the NaYF_4_:TmYb nanoparticles, with the Tm^3+^ transitions ^1^D_2_-^3^F_4_ and ^1^G_4_-^3^H_4_ noted. (c) Schematic representation of the upconversion process of Yb^3+^/Tm^3+^ codoped UCNPs. Blue luminescence is ascribed^14^ to ^1^D_2_-^3^F_4_ (blue arrow) and ^1^G_4_-^3^H_4_ (light blue arrow). The UV are ascribed to ^1^D_2_-^3^H_6_ (pink) and ^3^P_6_ – ^3^F_4_ (not shown).

In 2009 Branda and co-workers pioneered harnessing the output from UCNPs to activate (dithienylethene) photoswitches^18^. Subsequently, both spiropyran^19^ and AB^20^ photoswitches have been indirectly controlled using UCNP^21^. In parallel to these studies, biological photoswitches (light-gated ion channels^22–29^, and LOV domains^30^) have been stimulated by the luminescence from UCNPs. Here we show that blue luminescence generated by 975 nm excitation of a NaYF_4_:TmYb nanoparticle can be used to stimulate *cis* to *trans* isomerization of the mono quaternary ammonium photoprobe AAQ. Simple ABs such as AAQ are optically transparent to single photon excitation at 975 nm, and are poorly sensitive to direct 2-photon excitation^31^. In contrast, we found that when AAQ and UCNPs were co-loaded via a patch-clamp pipette into living neurons, we could use 975 nm light to trigger *cis* to *trans* isomerization of AAQ and block voltage-gated potassium channels in CA1 neurons in hippocampal brains slices.

## Results and discussion

We selected NaYF_4_:25%Yb,0.5%Tm nanoparticles, enveloped in PEG chains for aqueous compatibility, that were about 35 nm in size (Fig. 2a). When excited at 975 nm these luminescence mostly in the UV-blue range (Fig. 2b). The NPs show two narrow emission bands (Fig. 2b) that are characteristic of ^1^D_2_-^3^F_4_ (ca. 450 nm) and ^1^G_4_-^3^H_4_ (ca. 470 nm) transitions in Tm^3+^ activators (Fig. 2c). With additional bands in the near-UV, that are ascribed to ^1^D_2_-^3^H_6_ and ^3^P_6_ – ^3^F_4_ transitions^15^.

We dissolved these NPs in HEPES buffer at a concentration of 0.8 mg/mL with AAQ (0.040 mM) in a quartz cuvette (1 cm pathlength, volume 50 μL). The beam (diameter ca. 0.8 mm) from a Ti:sapphire laser (975 nm) produced blue luminescence that was easily seen with the naked eye at an intensity of 500 mW (Fig. 3a). Irradiation of all *trans*-AAQ with a light from a UV LED (365 nm) produced the *cis* photostationary state of AAQ (> 80% *cis* configuration^8^, Fig. 3c, dashed curve). *cis*-AAQ is quite unstable thermally, having a half-life of about 6 mins^8^. The rate of *cis* to *trans* isomerization can also be enhanced by irradiation with blue-green light^8^. Thus, after 30 s in the dark, the *cis*-AAQ absorption spectrum showed small changes due to some thermal relaxation (Fig. 3c, red curve). However, when irradiated with 975nm-light (1.2 W) for 30 s the absorption spectrum changed considerably (Fig. 3c, blue curve). On standing for a further 30 s at RT in the dark the shape of this spectrum had changed a little more (Fig. 3c, orange curve). These data suggest that blue light from NIR upconversion can significantly speed up the *cis* to *trans* isomerization of AAQ in the same way as blue/green light^8^. Encouraged by these chemical transformations observed in cuvette, we tested mixtures of NPs and AAQ in CA1 neurons in hippocampal brain slices acutely isolated from mice.

**Figure 3.**
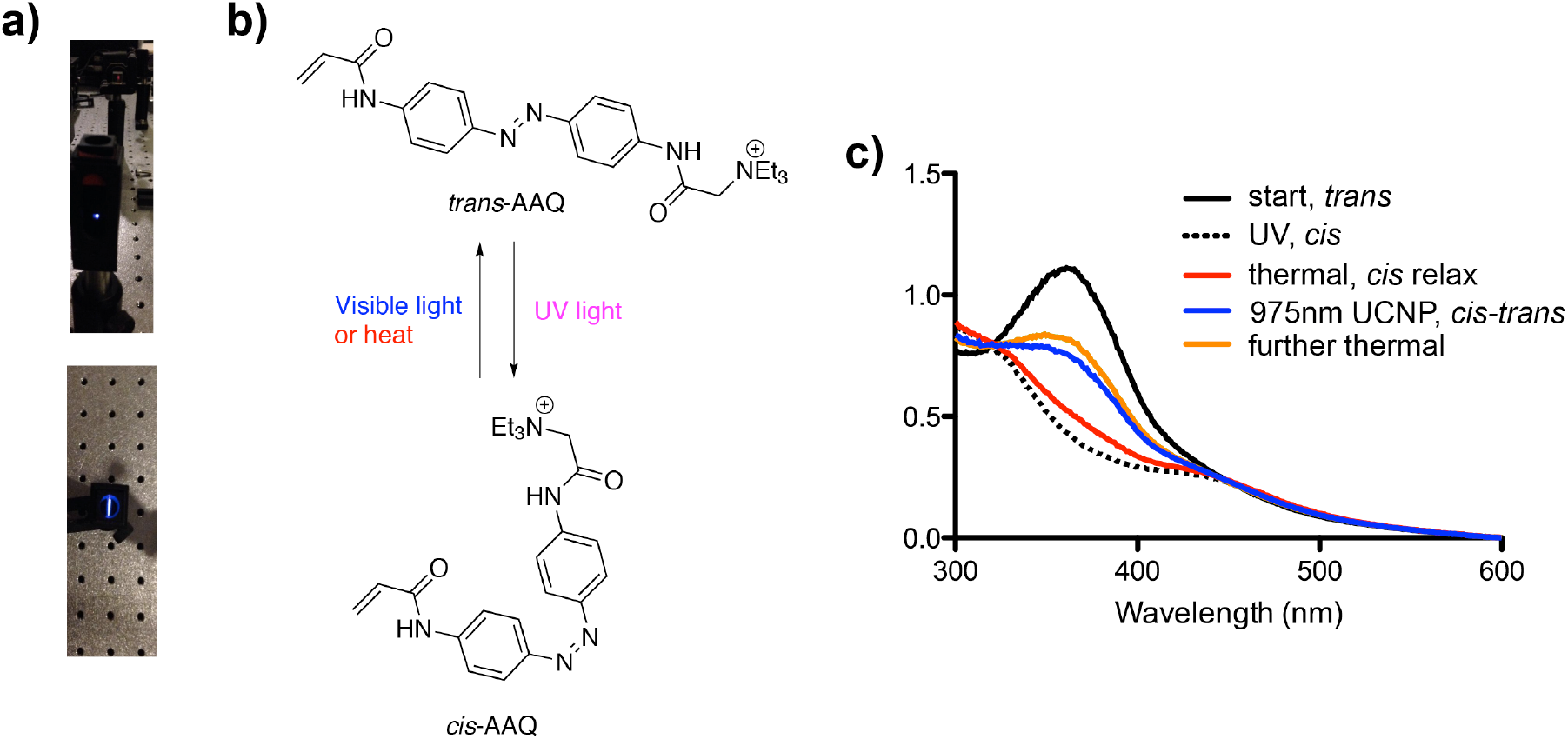
UCNP blue luminescence in cuvette and AAQ isomerization. The NaYF_4_:TmYb nanoparticles were dissolved with AAQ in HEPES buffer (pH 7.4). The solution (0.050 mL) was excited with a laser at 975 nm in a quartz cuvette (pathlength 1 cm). (a) Photographs of the blue luminescence from the side and top. (b) AAQ in *cis* and *trans* configurations. (c) UV-visible absorption spectrum of all *trans*-AAQ (black) and the UV photostationary state (i.e. *cis*) of AAQ (dashes). The red curve was taken following standing in the dark for 30 s at RT, the blue curve after 30 s irradiation with 975 nm light, and the orange curve after a further 30 s at RT in the dark.

First, we tested if the UCNPs themselves were toxic in pyramidal neurons. To do this we loaded NPs via a patch pipette into CA1 neurons at a concentration of 0.8 mg/mL. Neurons were held in the current clamp mode at −65 mV, and current injections lasting 200 ms were applied in steps of 20 pA from 0 to 180 pA. We performed this protocol at two time points: first after initial whole-cell access; then after 30 mins. We examined the threshold for firing action potentials, resting membrane potentials, action potential amplitudes and the spike rate in current clamp mode. Under voltage clamp mode, input resistance (R_in_) was measured and calculated. Voltage steps (ΔV, −20 mV for 100 ms) were given to the neurons and the peak steady-state current responses (ΔI) to the voltage steps were recorded. R_in_ was calculated as ΔV/ΔI. We could detect no statistically significant difference for these electrical properties between neurons containing or not containing the UCNPs (n = 3 cells, without and with UCNPs, Fig. 4). Our data are consistent with many reports that aqueous-compatible UCNPs are non-toxic towards living cells^14^, and are even well tolerated when implanted in living tissue over the long term^27,29^ (many weeks).

**Figure 4.**
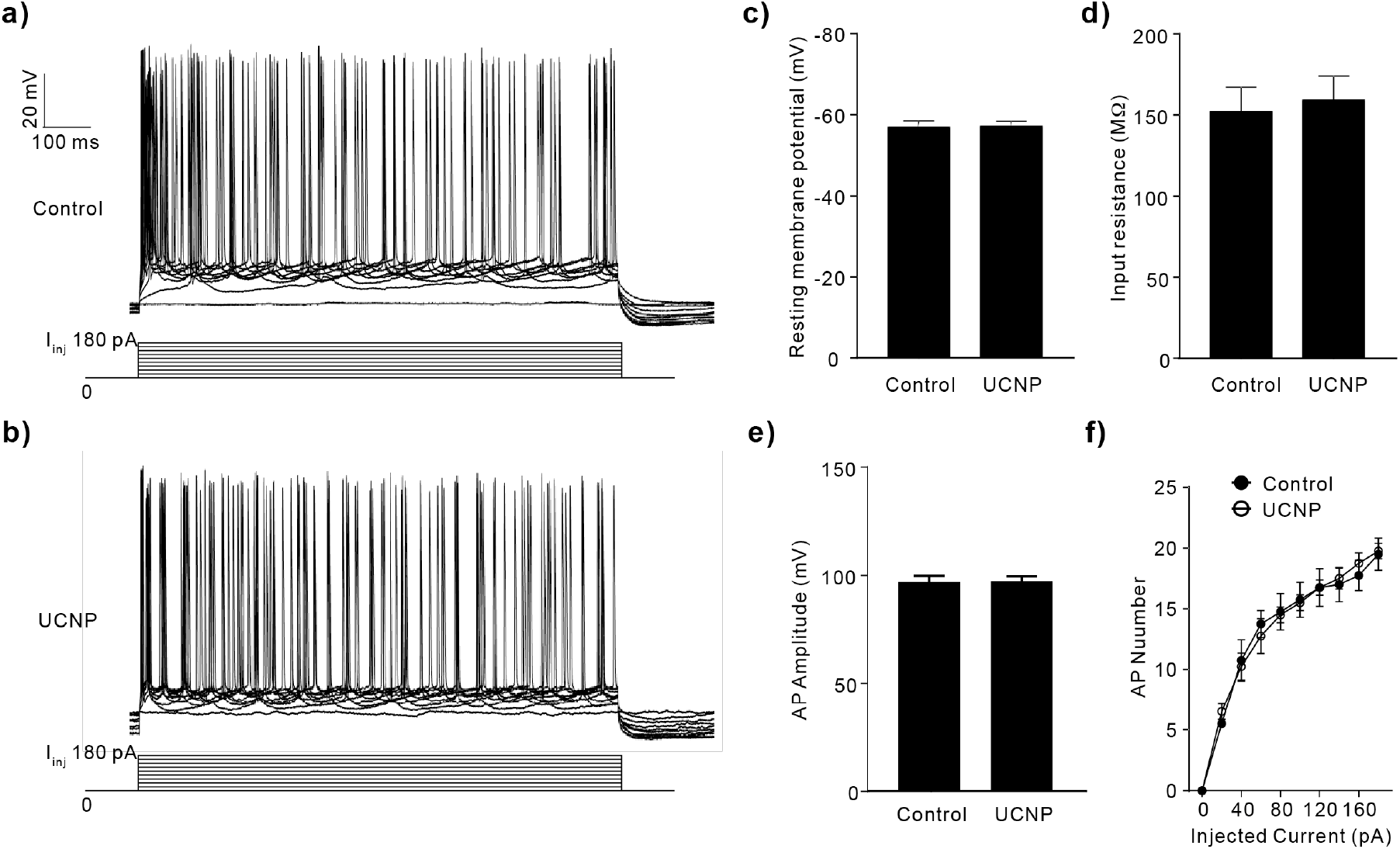
UCNPs are non-toxic inside cells. (a, b) Sample traces of current-clamp recordings of depolarizations and action potentials in CA1 pyramidal neurons during constant current injections (from 0 pA to 180 pA for 1 s). Internal recording solution used in control group was K-gluconate based, without or with 0.8 mg/mL UCNPs (n = 4 cells in each group). (c-e) Basic intrinsic properties of action potentials, including resting membrane potentials (c, t-test, two-tailed, t = 0.1122, df = 6, p = 0.9144), input resistance (d, t-test, two-tailed, t = 0.3305, df = 6, p = 0.7523), and action potential amplitudes (e, t-test, two-tailed, t = 0.1282, df = 6, p = 0.9022) were unchanged by UCNPs. (f) Numbers of action potentials induced by 1 s increasing current injections were unchanged by UCNPs (Two-way RM ANOVA, F _(9, 27)_ = 0.4475, p = 0.8963).

Even though NIR is considered less toxic to cells than UV or visible light, we needed to test what power level of NIR light CA1 neurons could tolerate. To do this we filled CA1 neurons via a patch pipette with UCNPs (0.8 mg/mL) along with the red fluorescent dye Alexa-594. This allowed us to image patch-clamped cells using two-photon laser-scanning microscopy at 810 nm. We then re-tuned the laser to 975 nm, and increased the pixel dwell time to 10 microseconds per pixel (Fig. 5b,c). Cells were scanned in contiguous 48 s epochs with increasing power. We found that between 55-60 mW some drift in baseline started to be detected by the electrophysiological probe. Powers above 60-65 mW caused cell death rapidly, whereas 30 and 40 mW were very well tolerated (n= 3 cells, Fig. 5a).

**Figure 5.**
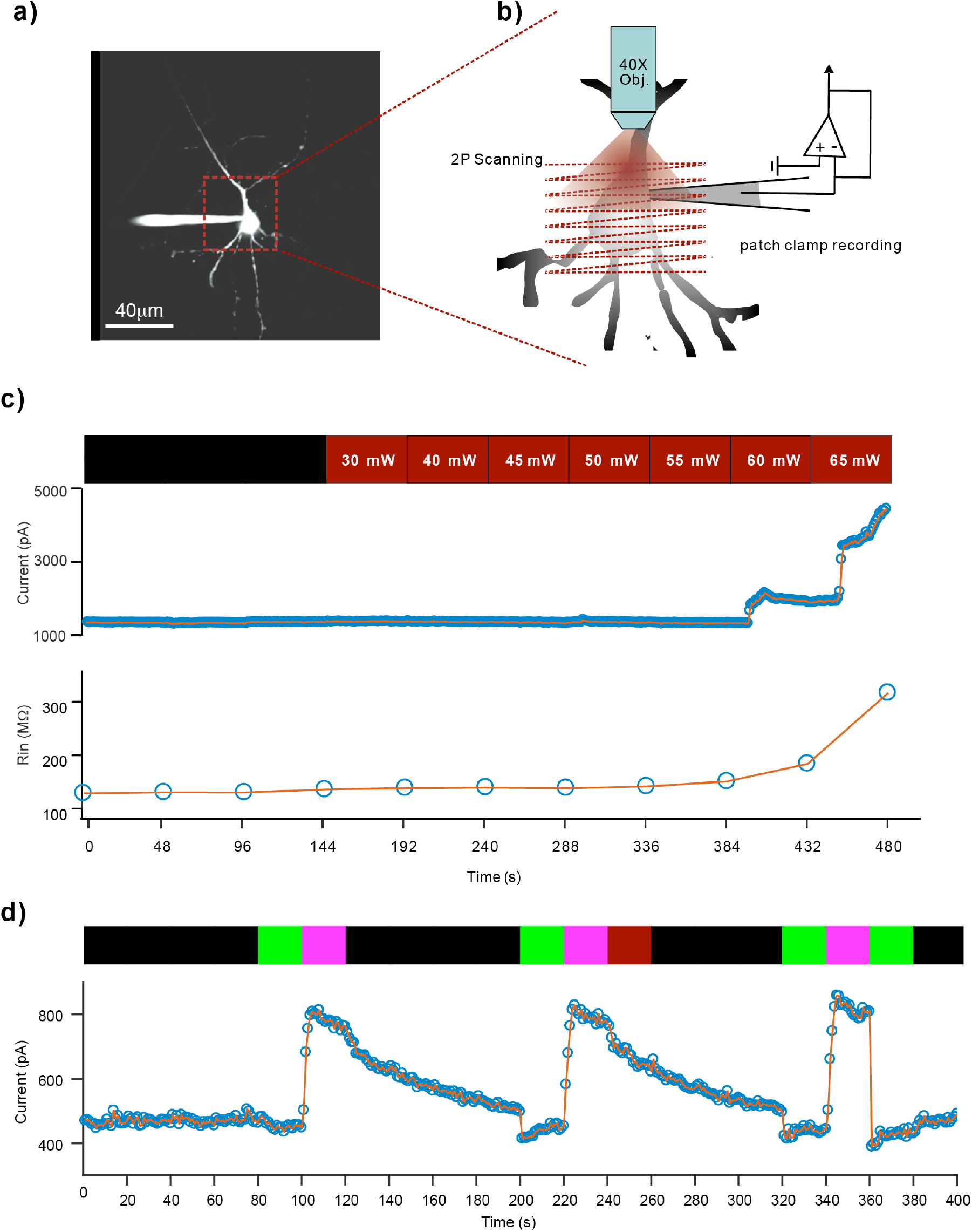
Power threshold for NIR irradiation of CA1 neurons. (a,b) Schematic diagram of NIR light irradiation of a CA1 neuron. A patch-clamped neuron was imaged at 810 nm using a two-photon microscope with a 40X objective (0.8 numerical aperture). The field of view was 171 microns. To induce photoswitching the soma of the neuron was scanned at 975 nm with a field of view of 64 microns with 10 μs dwell time. In both cases 512×512 pixels were used. (c) Top: Voltage-dependent currents generated by depolarizing voltage steps (−60 mV to +40 mV, 200 ms duration, 1Hz) were recorded from CA1 pyramidal neurons either with no light, or 975 nm illumination (powers 30, 40, 45, 50, 55, 60, 65 mW). Bottom: The input resistances of the cell during the recording were calculated in each 48-s epoch. The whole-cell internal solution was K-gluconate based with 0.4 mM AAQ. This test was repeated for 3 neurons. (d) Comparison the effect of 975 nm light on *cis* to *trans* isomerization of AAQ in neurons by measuring voltage-dependent currents. No light is indicated as black, UV LED irradiation (365 nm, 0.5 mW) as violet, green LED (530 nm, 1.5 mW) irradiation as green, and NIR light (975 nm, 30 mW) as dark red, each epoch lasted 20 sec. The test was repeated for 3 neurons.

Next we tested if NIR at low power (30 mW) had any effects on AAQ photopharmacology in CA1 neurons in the absence of UCNPs. To do this we filled CA1 neurons with Alexa-594 via a patch pipette, and imaged the cells using two-photon microscopy at 810 nm, as above. In this experiment we included AAQ in place of UCNPs as part of the internal solution. Quaternary ammonium ions are well known to block voltage-gated potassium ion channels from the inside of cells^32,33^. Such blockade is known to “use dependent”, meaning that at rest such organic cations do not have access to their internal binding site. However, voltage steps from −60 to + 40 mV, caused by patch-clamp control, allow organic cations access to the blocking site. In the case of AAQ it is has been determined that such steps must be applied at about 1 Hz^8^. In order to maintain this blockade such voltage steps must also be applied continuously, otherwise there is “unuse-dependent unblock”. We found that after establishing a good baseline with cells loaded with all *trans*-AAQ, irradiation of the cells with UV light (365 nm) induced rapid unblocking of voltage-gated potassium channels (initial black and violet epochs in Fig. 5d), as illustrated in Fig. 1. After 20 s of continuous illumination with 0.5 mW, irradiation was terminated, allowing slow increase of ion blockade over the next 80 s, due to the thermal *cis* to *trans* isomerization of AAQ. Irradiation with green light for 20 s reset blockade to pre-UV levels, which allowed us to repeat photoswitching with UV for an additional 20 s, causing a second period of ion channel unblock (Fig. 5b). Then we scanned the cell with NIR for 20 s (30 mW, as above), followed by an additional period of 60 s with no illumination at any wavelength. This second 80 s post-UV period was indistinguishable from the first when no NIR was applied. Finally, we applied green-UV-green illumination for 20 s epochs to confirm cell health and efficient AAQ photoswitching inside the cell. These data imply that the NIR power dosage we use had no effect on AAQ. Based on the control experiments in Fig. 5 we concluded the UCNPs could be used inside CA1 neurons in hippocampal brain slices with 30 mW of NIR light without any deleterious side effects.

Having established these basic parameters, we showed that blue luminescence from UCNPs could drive AAQ isomerization in CA1 so as to control voltage-gated potassium channels in situ. CA1 neurons were filled with UCNPs (0.8 mg/mL) along with the red fluorescent dye Alexa-594 and AAQ (0.4 mM) via a patch pipette. Again we established the baseline use-dependent blockade of potassium channels. After a period of a few mins, 20 s of green light (1.5 mW full field) caused a mild increase in blockade. Illumination with UV light for 20 s (0.5 mW full field) induced rapid unblock of potassium channels (Fig. 6), as above. A 20 s period in the dark induced a mild block, which was reversed by a further 20 s of UV light. Following this reset, we interleaved 20 s periods of NIR (30 mW, as above) scanning and UV illumination to cause bi-directional modulation of potassium channel blockade by AAQ. Finally, green light was applied to induce full blockade. Note that NIR was almost as effective as green light in this experiment, albeit at a slower rate (Fig. 6).

**Figure 6.**
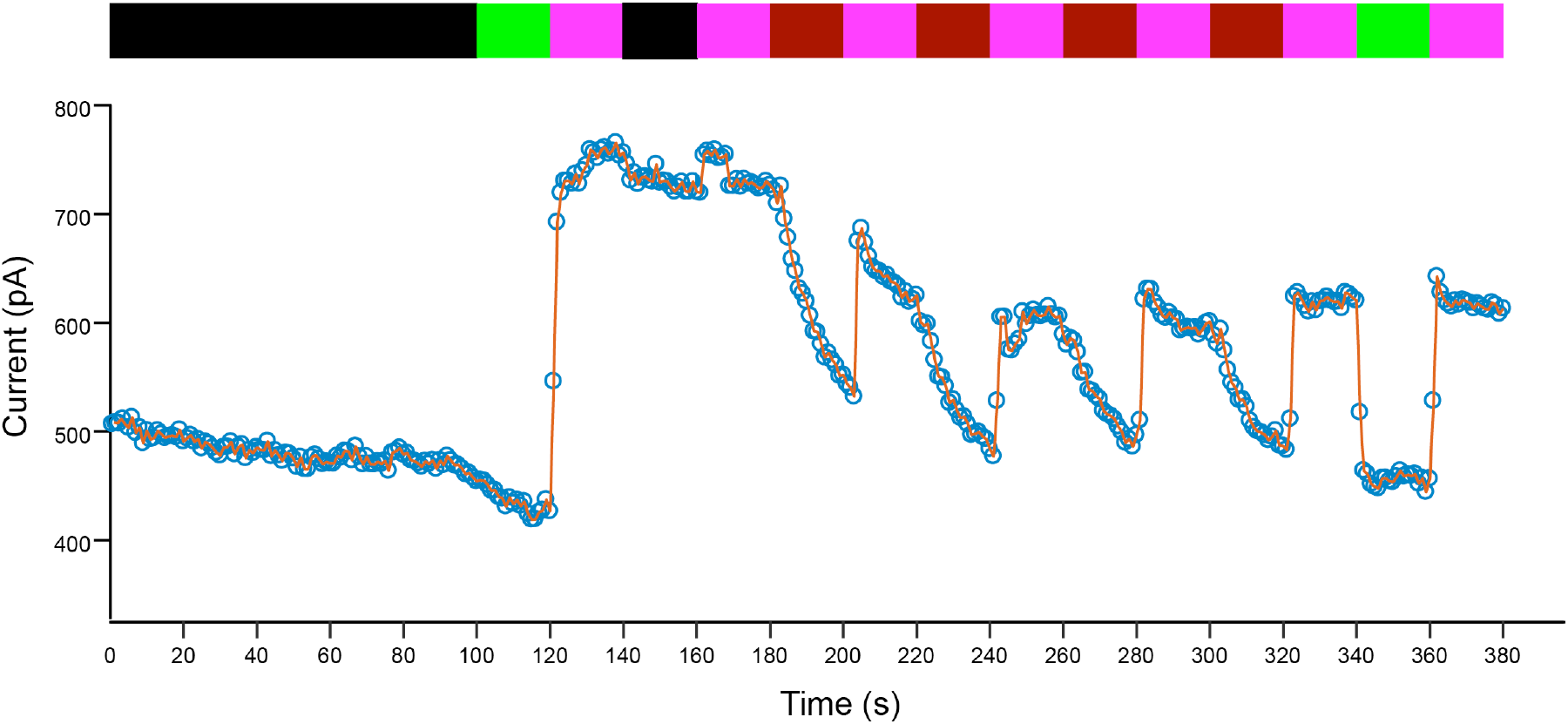
Bi-directional photoswitching of AAQ in CA1 neurons with UV and NIR light. Voltage steps (from −60 mV to + 40 mV for 200 ms) where applied continuously to CA1 neurons at a frequency of 1 Hz. Voltage-dependent currents at the end of the 200 ms period, which correspond to those produced from voltage-gated potassium channels are displayed during photoswitching of AAQ. The internal solution was K-gluconate based, with 0.4 mM AAQ and 0.8 mg/mL UCNPs. No light irradiation is indicated as black, UV LED irradiation (365 nm, 0.5 mW) as violet, green LED (530nm, 1.5 mW) irradiation as green, and NIR (975 nm, 30 mW) as dark red, each epoc lasted 20 s. The test was repeated in 3 different CA1 neurons.

## Conclusions

Use of UCNPs to control neuronal membrane potential is a topic of current interest^34^. Here we show that these materials can also be co-loaded with a photoswitchable drug into living neurons via a patch pipette, and used to control ion channel conductivity. This approach is complementary to those proposed in other studies, with some encasing UCNPs in polymer films placed adjacent to opsin-transfected neurons^22–24^, linking UCNPs intimately to ion channels^25,26^, or simply injecting UCNPs into the extracellular space of living mice^27,29^. Our study relies on the physical confinement provided by the intracellular environment to bring the UCNPs into intimate proximity with an azobenzene (AB) photoswitch called AAQ^8^ in CA1 neurons in living brain slices. Co-loading neurons with both chromophores allows the blue luminescence from the UCNPs to be re-absorbed by AAQ and drive the *cis* to *trans* isomerization that is crucial to control of the AAQ photoprobe^8^. The chromophore used in AAQ is common to that in many other AB photoswitches^3^, thus our work suggests that upconversion could be useful for “remote control” of such probes in complex biological systems using NIR light.

## Materials and methods

All chemical were obtained from commercial sources and used without further purification. AAQ was from Tocris Bioscience (Bristol, UK), and the NaYF_4_:TmYb nanoparticles from Creative Diagnostics (cat. no. dni-f001, Shirley, NY, USA).

UV-Vis absorption spectra were recorded using a Cary 50 spectrophotometer (Agilent, Santa Clara, CA, USA). Photolysis used UV or green LED (M365LP1, M530L3, Thorlabs, NJ, USA), and Ultra II laser (Coherent, Palo Alto, CA, USA)

All animal studies were approved by Mount Sinai IACUC review. C57BL/6J mice (8-12 weeks.) were anaesthetized with isoflurane and the brain was quickly removed. Horizontal slice sections (350 μm) were then made in ice-cold cutting solution containing (in mM): 60 NaCl, 2.5 KCl, 1.25 NaH_2_PO_4_, 7 MgCl_2_, 0.5 CaCl_2_, 26 NaHCO_3_, 10 glucose, 100 sucrose, 3 sodium pyruvate, 1.3 sodium ascorbate equilibrated with 95% O_2_ and 5% CO_2_ (pH 7.3-7.4). The brain slices were then incubated for 15 min at 33°C in artificial cerebrospinal fluid (ACSF, mM: 125 NaCl, 2.5 KCl, 1.25 NaH_2_PO_4_, 1 MgCl_2_, 2 CaCl_2_, 26 NaHCO_3_, 10 glucose, 3 sodium pyruvate, 1.3 sodium ascorbate; 95% O_2_ and 5% CO_2_, pH 7.3–7.4). After incubation, all brain slices were held in ACSF at room temperature > 1 hour before patch clamp recording.

Brain slices were transferred to a submerged chamber on a BX-61 microscope (Olympus, Penn Valley, PA, USA) and perfused with ACSF (in mM: 125 NaCl, 2.5 KCl, 1.25 NaH_2_PO_4_, 1 MgCl_2_, 2 CaCl_2_, 26 NaHCO_3_, 10 glucose, 95% O_2_ and 5% CO_2_, pH 7.3–7.4) at room temperature. Hippocampal CA1 pyramidal neurons were visualized with a 40X objective (Olympus) and infrared differential interference contrast optics. Whole-cell recordings were made with an EPC-10 amplifier (HEKA Instruments, Bellmore, NY, USA) in voltage-clamp mode (Vm holding at −60 mV) or current-clamp mode. The electrical signals were recorded at 20 kHz and filtered at 3 kHz with Patchmaster (HEKA). Recording pipettes (3-5 MΩ) were filled with an internal solution (in mM): 135 potassium gluconate, 4 MgCl_2_, 10 HEPES, 4 Na_2_-ATP, 0.4 Na_2_-GTP, 10 Na_2_-phosphocreatine, (pH 7.35). Glutamate receptor antagonists CNQX (10 μM) and DL-AP5 (100 μM) and the GABA_A_ receptor antagonist bicuculline (20 μM) were added to ACSF during experiments.

UV and green LEDs (365 and 530 nm, respectively) were mounted on the epifluorescence port of the microscope and light beams were combined and directed to the objective by standard long-pass dichroic mirrors. Light power was controlled via the LED driver (LEDD1B, Thorlabs) by external voltage modulation. Timing and duration of LED lights were controlled by TTL signals and synchronized with EPC-10 system. NIR excitation used 975 nm output from a Vision II laser, directed onto the patch-clamped cell via an Ultima scan head (Prairie Technologies, Middleton, WI, USA).

## Acknowledgements

This work was supported by grants from the NIH (USA) to GCRE-D.

## Author contributions.

JZ conducted all the biological studies. GCRE-D performed the cuvette experiments. GCRE-D supervised the work and wrote the paper. All authors approved the final submitted version.

